# Multifunctional Roles of Brr6 and Brl1 in Nuclear Envelope Fusion During Nuclear Pore Complex Biogenesis

**DOI:** 10.1101/2025.07.22.665954

**Authors:** Sayan Mondal, Annett Neuner, Azqa Khan, Jlenia Vitale, Elmar Schiebel

## Abstract

Brl1 and Brr6 are essential, paralogous integral membrane proteins of the yeast nuclear envelope (NE) that transiently associate with nuclear pore complexes (NPCs) during their assembly to promote fusion of the inner (INM) and outer nuclear membranes (ONM). An amphipathic α-helix (AαH) in Brl1 is critical for mediating this fusion during NPC biogenesis. However, the exact roles of Brl1 and Brr6 in the molecular mechanisms of NPC assembly are still unclear. Here, we demonstrate that Brr6 operates at both early and late stages of NPC assembly. Its early function is supported by AαH mutants that fail to permit nucleoporin recruitment and INM deformation, while mutations in conserved cysteine residues lead to NE herniations and defective membrane fusion. Additionally, the N-terminus of Brl1 interacts with Nic96, likely promoting its recruitment to nascent NPC assembly sites. We further provide evidence that the length of the perinuclear space-spanning region of Brl1 and Brr6 is critical for proper NE fusion during NPC formation. Artificial elongation of this region produces a toxic phenotype marked by Nup82 mislocalisation and severe NE integrity defects. This phenotype supports a model in which Brl1 and Brr6 promote NE fusion following INM deformation, when the distance between the INM and ONM is reduced—a process initiated by nucleoporin assembly on the nuclear side of the INM.

## Introduction

In eukaryotic cells, the genome is encapsuled by the nuclear envelope (NE), a spherical double membrane (Lin & Hoelz, 2019; Strambio-De-Castillia *et al*, 2010). Transport of macromolecules, RNA and proteins, in and out of the nucleus, is bidirectional and facilitated by nuclear pore complexes (NPCs) that are embedded into fusion sites of the outer (ONM) and inner nuclear membrane (INM) (Beck & Hurt, 2017). NPCs consisting of multiple copies of 30 different nucleoporins (Nups) that can be divided into four categories. Transmembrane Nups form a ring structure that anchors the NPC to the nuclear envelope (NE), while the inner and outer ring Nups constitute the core scaffold of the NPC, which also contributes to the formation of the central transport channel (Strambio-De-Castillia *et al*., 2010). Phe-Gly (FG) Nups are anchored to the core scaffold by linker Nups and are responsible for the selective permeability barrier of NPCs (Beck & Hurt, 2017; Frey & Gorlich, 2007).

Despite the essential functions of the NPC, the mechanisms underlying its assembly remain poorly understood. In higher eukaryotes, NPCs assemble via two distinct pathways. During open mitosis, the NE breaks down in prophase, leading to NPC disassembly. Following chromosome segregation, the NE and NPCs must be reassembled during mitotic exit, marking the transition from mitosis to G1 phase. In addition to this post-mitotic assembly, NPCs also form during interphase through an inside-out mechanism. In this pathway, Nups first accumulate on the nuclear side of the INM followed by local INM deformation, fusion of the INM and ONM, and eventual insertion of the assembled NPC into the NE (Otsuka *et al*, 2016; Otsuka & Ellenberg, 2018; Otsuka *et al*, 2018). Some cytoplasmic Nups attach to the NPC after its embedding into the NE.

Because the budding yeast *Saccharomyces cerevisiae* undergoes closed mitosis, where the NE remains intact throughout cell division, NPCs assemble exclusively via the interphase pathway (Winey *et al*, 1997). Yeast NPC assembly also follows an inside-out mechanism, as evidenced by mutations in some genes encoding Nups such as *NUP116* or involved in NPC biogenesis such as *BRL1* and *BRR6* (de Bruyn Kops & Guthrie, 2001; Onischenko *et al*, 2017; Wente & Blobel, 1993; Zhang *et al*, 2018). These mutations lead to herniations, deformations of the INM that protrude into the perinuclear space and contain Nups on the nuclear-facing side of the INM. Herniations probably arise because NPC assembly is successfully initiated at the INM but fusion between the two nuclear membranes fails (Onischenko *et al*., 2017; Thaller & Lusk, 2018; Zhang *et al*., 2018).

The order of Nup incorporation into the emerging yeast NPC was recently analyzed by metabolic pulse labeling with heavy isotopes followed by mass spectrometry analysis of the newly formed Nup subcomplex, an approach that was named KARMA (Onischenko *et al*, 2020). This indicates that the early tier group containing Nic96, Nup53, Nup57 and Nup84 become incorporated first, followed by intermediate (Nup85, Nup159 and Nup1 groups) and late tiers (Mlp1 and Mlp2) (Onischenko *et al*., 2020).

In *Saccharomyces cerevisiae*, the interacting, integral membrane proteins Brl1, Brr6 and Apq12 are important for assembly of new NPCs without being components of fully assembled NPCs (Hodge *et al*, 2010; Lone *et al*, 2015; Scarcelli *et al*, 2007; Zhang *et al*, 2021; Zhang *et al*., 2018). Apq12, Brl1 and Brr6 each contain two transmembrane regions (TM), which are connected by a stretch of amino acids that localize in the perinuclear space (Vitale *et al*, 2022; Zhang *et al*., 2021; Zhang *et al*., 2018). The perinuclear amino acids of Apq12 fold into a short amphipathic α-helix (AαH) that was shown to coordinate Brl1-Brr6 interaction and regulate lipid composition (Zhang *et al*., 2021).

In Brl1 and Brr6 the perinuclear stretch also carries an AαH and an additional disulfide-stabilized anti-parallel helix bundle (DAH). Several observations indicate a late role of Brl1 during NPC assembly namely in the fusion of the INM with the ONM and therefore in the embedding of the emerging NPC into the NE. First, depletion of Brl1 results in the formation of herniations (Zhang *et al*., 2018). Second, overexpression of *BRL1* mutants that impair AαH function, trigger the formation of INM petals with attached NPCs on its base suggesting continues expansion of the INM without fusion to the ONM (Kralt *et al*, 2022; Vitale *et al*., 2022). Finally, mild overexpression of *BRL1* suppresses the herniation defect in *nup116Δ* cells meaning that it enables fusion between the INM and ONM (Vitale *et al*., 2022; Wente & Blobel, 1993; Zhang *et al*., 2018).

It has been suggested that Brl1 and Brr6 carry out similar functions, but at different sides of the NE, as Brr6 localizes to the INM and ONM, while Brl1 is predominately detected at the INM (Kralt *et al*., 2022; Vitale *et al*., 2022; Zhang *et al*., 2018). Brl1 and Brr6 could connect the INM with the ONM as soon as the INM was bend to the nuclear side of the INM due to the deposition of Nups (Kralt *et al*., 2022; Vitale *et al*., 2022). However, the specific role of Brr6 has not yet been investigated, and it remains unclear why it associates with both leaflets of the NE, potentially indicating a dual role for Brr6 in NPC assembly (Zhang *et al*., 2018).

In this study, we investigate the functions of Brl1 and Brr6 in NPC biogenesis. Surprisingly, overexpression of *BRR6* mutants affecting the integrity of the AαH, specifically *brr6^L145E^* and *brr6^F152E^*, resulted in a strikingly different phenotype compared to analogous *brl1^F391E^* mutations, which caused the accumulation of large petal-like structures with defective INM/ONM fusion (Kralt *et al*., 2022; Vitale *et al*., 2022). Overexpression of *brr6^L145E^* and *brr6^F152E^*, as well as incubation of the cold-sensitive *brr6^L145E^* mutant at 16°C, disrupted NPC assembly at an early stage, without detectable INM deformation or Nup enrichment at the NE. By comparison, temperature sensitive *brr6(ts)* mutants, affecting the function of the DAH incubated at 37°C exhibited a herniation phenotype, indicative of an INM/ONM fusion defect. We also found that the essential N-terminus of Brl1 interacts with the Nic96 complex likely contributing to the recruitment of Brl1 to assembling NPCs. Notably, elongation of the DAH domain in either Brl1 or Brr6 impaired function; its overexpression was lethal and triggered partially disintegration of the NE. Taken together, these findings indicate that Brl1 and Brr6 perform multiple essential roles in NPC biogenesis and support a model in which the length of the DAH is a critical determinant for proper NE fusion during NPC assembly.

## Results

### AlphaFold predictions of Brl1 and Brr6 interactions

Previous studies have shown that Brl1 and Brr6 interact physically (Lone *et al*., 2015; Zhang *et al*., 2021; Zhang *et al*., 2018). Brl1 is predominantly localized to the INM, whereas Brr6 exhibits dual localization at both the ONM and INM (Zhang *et al*., 2018). These distribution patterns suggest the potential for multiple modes of interaction between Brl1 and Brr6. To explore this, AlphaFold modeling (Jumper *et al*, 2021) of Brl1 and Brr6 (Figure 1a, b) was employed, which predicted an association mediated by their two AαHs (Figure 1c). Additionally, AlphaFold suggested head-to-head interactions between Brl1 and Brr6 involving the tip domains of their DAHs (Figure 1d).

**Figure 1.**
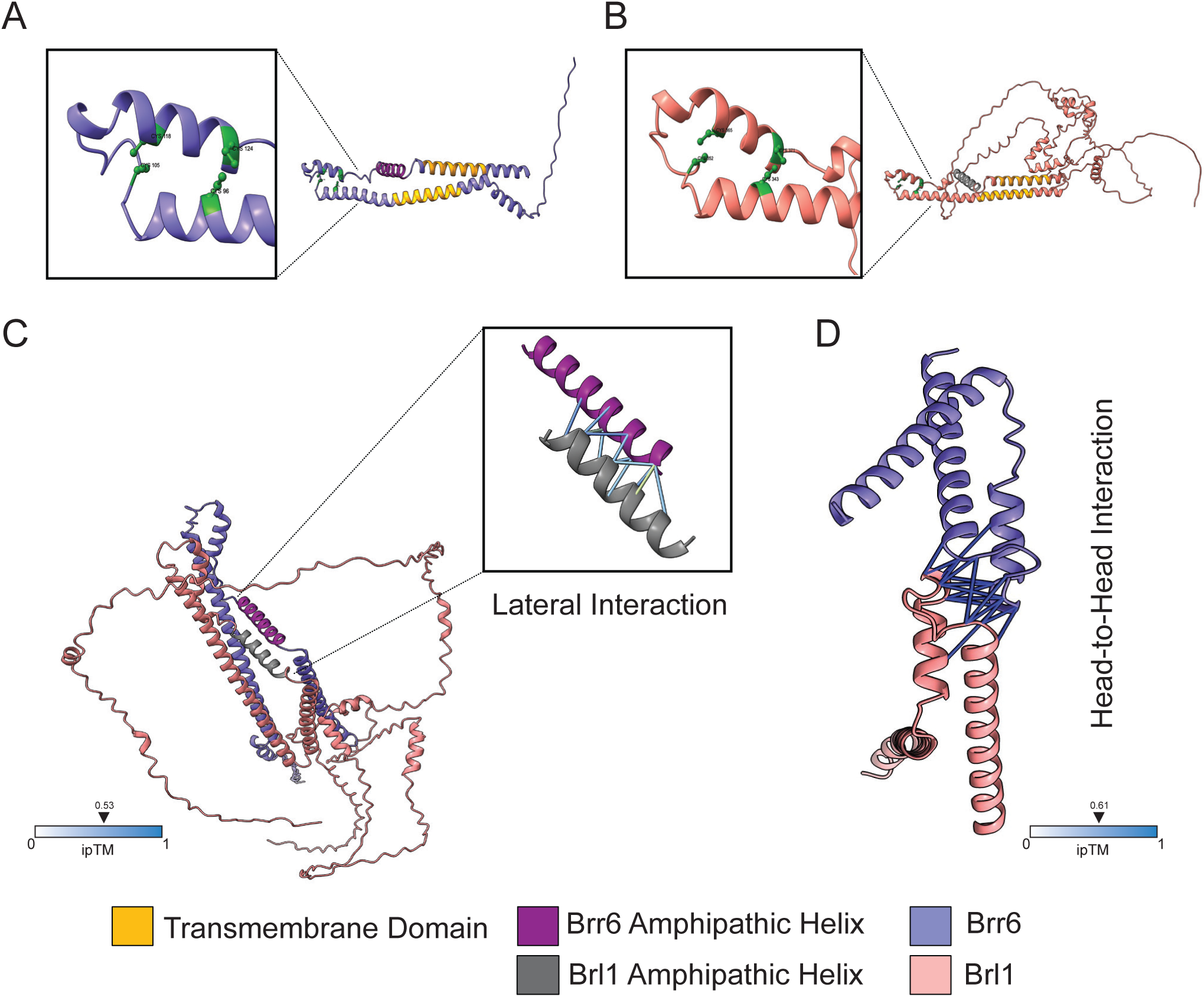
AlphaFold predictions of Brl1 and Brr6 interactions. **(A)** Domain organization of Brr6. An AαH is flanked by two transmembrane (TM) domains. Two disulfide bonds are indicated. **(B)** Domain organization of Brl1. The amphipathic helix is in dark grey and the four cysteine residues are also shown in green. **(C)** AlphaFold2 predictions of full length Brr6 and Brl1 with the interaction surfaces between their AαHs. **(D)** The interaction surface between the head-to-head domains of both Brr6 and Brl1 predicted using AlphaFold2.

The lateral AαHs may mediate interactions at the INM where both proteins localized (Zhang *et al*., 2018). The DAH tip-to-tip interactions could facilitate contacts across the NE, bridging Brl1 on the INM and Brr6 on the ONM through the intermembrane space.

### AαH of Brr6 performs an essential function required for yeast viability

The functional importance of the AαH in Brl1 for INM–ONM fusion during NPC assembly was demonstrated by the defective attempts of fusion of the deformed INM with the ONM in overexpressed *BRL1* AαH mutants (Kralt *et al*., 2022; Vitale *et al*., 2022). We addressed the function of the AαH in Brr6 following a similar strategy by introducing the AαH disrupting mutations L145E and F152E in *BRR6* (Figure 2a). We first tested whether the F152E mutation affects the ability of the AαH to associate with membranes. To do this, we expressed GFP-tagged fragments corresponding to the wild-type AαH (*BRR6^AαH^-WT-GFP*) and the mutant AαH (*brr6^AαH-F152E^-GFP*) in yeast. *BRR6^AαH-WT^*-GFP localized predominantly to the NE and the cytoplasmic membrane, both of which were marked by dsRED-HDEL (Munro & Pelham, 1987) (Figure 2b). In contrast, brr6^AαH-F152E^-GFP showed only weak NE localization (Figure 2b). These findings indicate that membrane interaction by the AαH of Brr6 depends on its amphipathic character.

**Figure 2.**
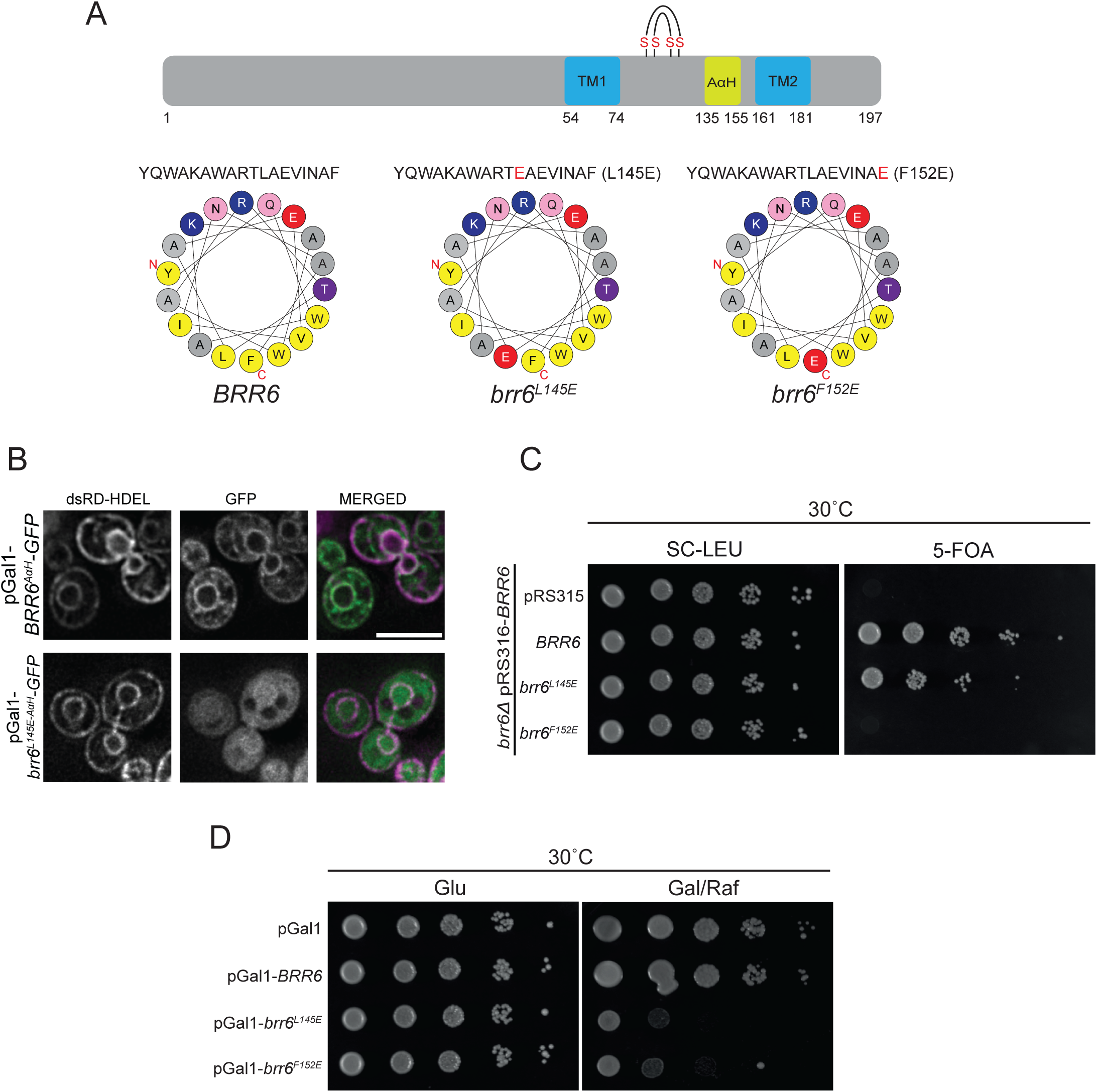
AαH of Brr6 performs an essential function required for yeast viability. **(A)** Amphipathic helix predictions for the wild-type and mutant sequences of Brr6 were generated using HeliQuest. The upper cartoon illustrates the domain organization of Brr6, highlighting the two transmembrane (TM) domains, the AαH, and the four conserved cysteine residues. Amino acid positions are indicated. **(B)** Subcellular localization of GFP-tagged Brr6 wild-type and brr6^L145E^ AαH region. Fluorescence microscopy was performed to assess localization. Scale bar: 5 µm. **(C)** Growth assay of the *BRR6* shuffle strain carrying the empty *LEU2*-based plasmid pRS315 or pRS315 containing the indicated *BRR6* alleles (*BRR6*, *brr6^L145E^*, and *brr6^F152E^*). Tenfold serial dilutions were spotted onto SC–LEU and 5-FOA plates and incubated at 30°C. **(D)** Wild-type yeast cells carrying the indicated pGal1 plasmids were spotted in tenfold serial dilutions onto glucose (Glu) or galactose/raffinose (Gal/Raf) plates and incubated at 30°C.

To test the functional importance of the AαH in Brr6, we examined the effects of the *brr6^L145E^* and *brr6^F152E^* mutations using a plasmid shuffle approach. Cells with *brr6^F152E^* failed to grow, indicating that the integrity of the AαH is essential for Brr6 function (Figure 2c). Cells expressing *brr6^L145E^* formed colonies with reduced efficiency (Figure 2c); and *brr6^L145E^*cells exhibited cold sensitivity (Figure 4a).

We next assessed the impact of overexpressing *brr6^L145E^*and *brr6^F152E^* on yeast growth. Cells overexpressing these mutant alleles displayed severe growth defects on the inducing galactose plates while control cells (pGal1-*BRR6* and pGal1 vector) showed robust growth (Figure 2d). In summary, the AαH of Brr6 performs an essential function required for yeast viability.

### The AαH of Brr6 is needed early in NPC assembly

We tested if overexpression of *brr6^L145E^* and *brr6^F152E^*affects NPC biogenesis. GFP-tagged Nups associated with different NPC substructures were analyzed in this experiment (Figure 3a, Supplementary Fig. 1). The controls, pGal1 plasmid or overexpression of *BRR6*, did not impact Nup localization along the NE (Figure 3a, b). In contrast, *brr6^L145E^* and *brr6^F152E^* over-expression impaired NE localization of most Nups, which accumulated in dots that localized close to the NE (Figure 3a, Supplementary Figure 1a-c). Line scan analysis of the Nup-GFP signals along the NE (Supplementary Figure 1b), allowed us to group the Nups into three classes (Figure 3b). Those most heavily affected, with approximately 50-60% mislocalization (Group 1); intermediately affected Nups (Group 2; 20-45%); and Nups that retained NE localization (Group 3; Figure 3b). Interestingly, Nic96 localization was intermediately affected by approximately 40–45% upon overexpression of *brr6^L145E^*and *brr6^F152E^* (Figure 3b). Transmembrane Nups, Ndc1 and Pom152, and the nuclear FG repeat Nup2 belonged to the group 3 category (Figure 3b).

**Figure 3.**
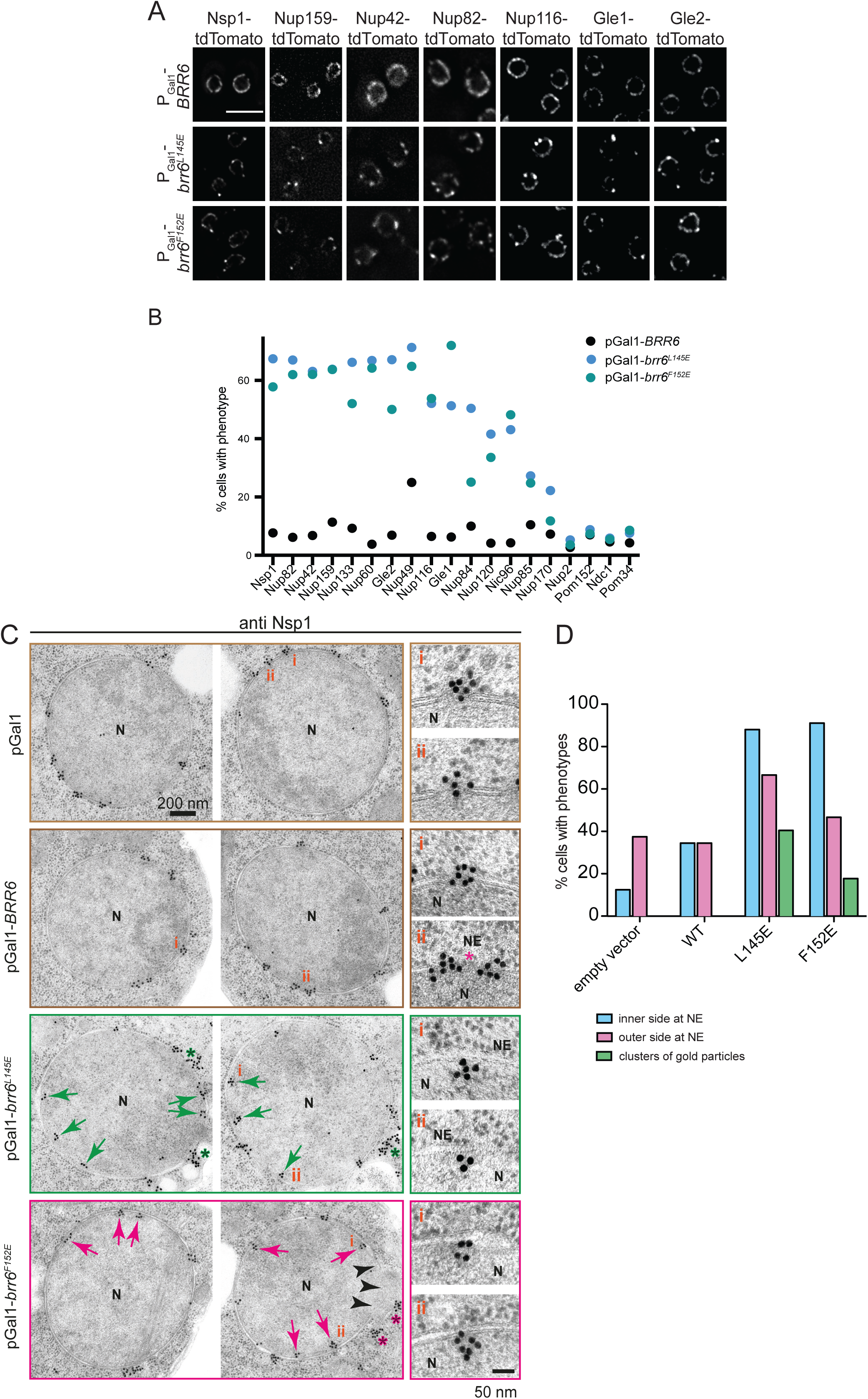
The AαH of Brr6 is needed early in NPC assembly. **(A)** Overexpression of *brr6^L145E^* alters NE localization of Nups. *BRR6*, *brr6^L145E^* and *brr6^F152E^* were overexpressed for 3 hours in yeast cells expressing fluorescently tagged (tdTomato) components of the NPC. Scale bar: 5 µm. **(B)** Quantification of the mislocalization phenotype across different tagged Nups. The graph shows the percentage of cells exhibiting mislocalization upon overexpression. N = 3 independent experiments. **(C)** Immuno-electron micrographs of wild-type cells overexpressing the control plasmid, *BRR6*, *brr6^L145E^*, or *brr6^F152E^* for 3 hours, stained with anti-Nsp1 antibodies. Red and green arrows indicate Nsp1 signal below the INM. Red asterisk highlight clusters of gold particles. Scale bar: 200 nm. Enlarged views of selected regions are shown to the right. Scale bar: 50 nm. **(D)** Quantification of gold particle labeling from (C). Clusters containing ≥10 gold particles were counted separately, as these likely correspond to the intense Nsp1 fluorescence foci observed in Supplementary Figure 1.

We next analyzed the phenotypes of *brr6^L145E^* and *brr6^F152E^* overexpression by EM. Surprisingly, most cells overexpressing pGal1-*brr6^L145E^* or pGal1-*brr6^F152E^*did not accumulate petal-like structures or herniations as this was seen in pGal1-*brl1^AαH^* or *brl1* mutant cells (Kralt *et al*., 2022; Vitale *et al*., 2022; Zhang *et al*., 2018) (Figure 3c). Immuno-EM analysis of the nucleoporin Nsp1 localization (Zhang *et al*., 2018) in cells overexpressing *pGal1-brr6^L145E^* and *pGal1-brr6^F152E^* revealed Nsp1 signals near the INM that were not associated with NPCs. This contrasts with control cells, where Nsp1 was mainly associated with NPCs (Figure 3c, enlargement right). In 20-40% of pGal1- *brr6^L145E^* and pGal1-*brr6^F152E^* cells, the Nsp1 signal was clustered on the cytoplasmic side of the NE as indicated by >10 gold particles (Figure 3c, d). This together suggests that overexpression of *brr6* mutants with a defective AαH impairs NPC assembly at an early stage before deformation of the INM.

### Cold sensitive *brr6^L145E^* cells are devoid of NPCs

The *brr6^L145E^* mutant cells exhibited cold sensitivity for growth (Figure 4a). At 16°C, analysis using Pom152 and Nup159 as NPC markers revealed a localization defect for Nup159, but not for Pom152 (Figure 4b, c), similar to pGal1-*brr6^L145E^* and pGal1- *brr6^F152E^* cells (Figure 3). EM analysis showed that the number of NPCs per NE section decreased from four in control cells to basically zero in *brr6^L145E^* cells (Figure 4d, e). Notably, the NE in *brr6^L145E^* cells incubated at 16 °C showed only a small number of herniations (Figure 4d, f). However, we noticed accumulation of ER-like structures in the cytoplasm of *brr6^L145E^* cells incubated at 16°C that contained elongated electron dense sections of disrupted membranes (Figure 4c, bottom and Figure 4f). These electron dense regions were labelled by the Nsp1 antibodies (Figure 4c, right, red asterisk). This phenotype suggests that because *brr6^L145E^* cells cannot assemble NPCs in the NE, Nup containing assemblies accumulate in the ER.

**Figure 4.**
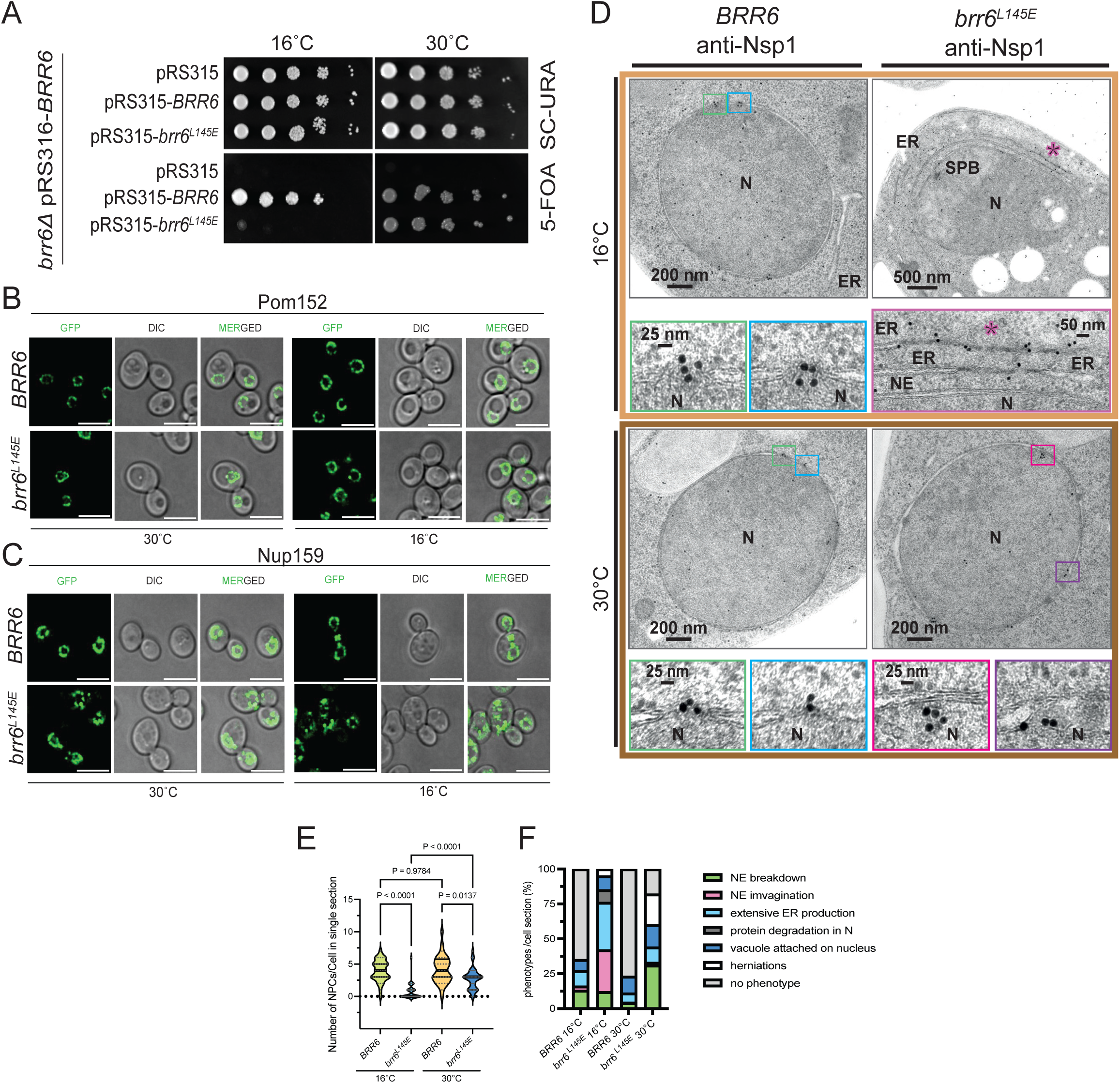
The AαH of Brr6 is needed early in NPC assembly. **(A)** The *brr6^L145E^* mutant exhibits cold-sensitive growth. To assess functionality at 16°C, Δ*brr6* cells carrying the *URA3*-based plasmid pRS316-*BRR6* were transformed with the indicated *LEU2*-based plasmids. Transformants were grown at both 16°C and 30°C. Growth on 5-FOA plates selects for cells that have lost the *URA3* plasmid, thereby testing whether the *LEU2*-based plasmid alone can support viability in the absence of wild-type *BRR6*. **(B, C)** Phenotypic analysis of *brr6^L145E^* cells. The *brr6^L145E^*allele was genomically integrated into Δ*brr6* pRS316-*BRR6* cells, followed by counterselection on 5-FOA. *POM152* (B) and *NUP159* (C) were tagged with GFP using genomic integration (Janke *et al*, 2004). Cold-sensitive *brr6^L145E^* cells were incubated at 16°C overnight and imaged by fluorescence microscopy. Scale bars: 5 µm. **(D)** *brr6^L145E^* cells analyzed by immuno-electron microscopy using anti-Nsp1 antibodies. In wild-type cells, Nsp1 was predominantly detected at the NPCs. In *brr6^L145E^* mutant cells, the Nsp1 signal was observed below the INM. Additionally, *brr6^L145E^* cells accumulated electron-dense regions within the ER that were specifically labeled by Nsp1 antibodies. **(E)** Quantification of NPCs per EM section from (D). *BRR6* (16°C): N = 31; *BRR6* (30°C): N = 31; *brr6^L145E^* (16°C): N = 32; *brr6^L145E^* (30°C): N = 35, sections were analyzed per strain and condition. Statistical analysis: ANOVA. (**F)** Quantification of phenotypes observed in (D). Cells were categorized as outlined in the figure. *BRR6* (16°C): N = 57; BRR6 (30°C): N = 77; *brr6^L145E^* (16°C): N = 78; *brr6^L145E^* (30°C): N = 67.

Taken together, these results support conclusions drawn from overexpression studies of *brr6* AαH mutants and indicate that Brr6 has a function early in NPC assembly, which requires a functional AαH.

**Brl1 and Brr6 AαH domain mutants show cross complementation** Overexpression of AαH domain mutants in *BRL1* and *BBR6* result in strikingly different phenotypes. We therefore analyzed the phenotype of *brl1^F391E^ brr6^L145E^* co-overexpression in order to understand the cooperativity of their interaction. Interestingly, co-overexpression of pGal1-*brl1^F391E^* pGal1-*brr6^L145E^* rescued the toxic effect of expression of the single mutants (Figure 5a) suggesting cross-complementation between both mutants.

**Figure 5.**
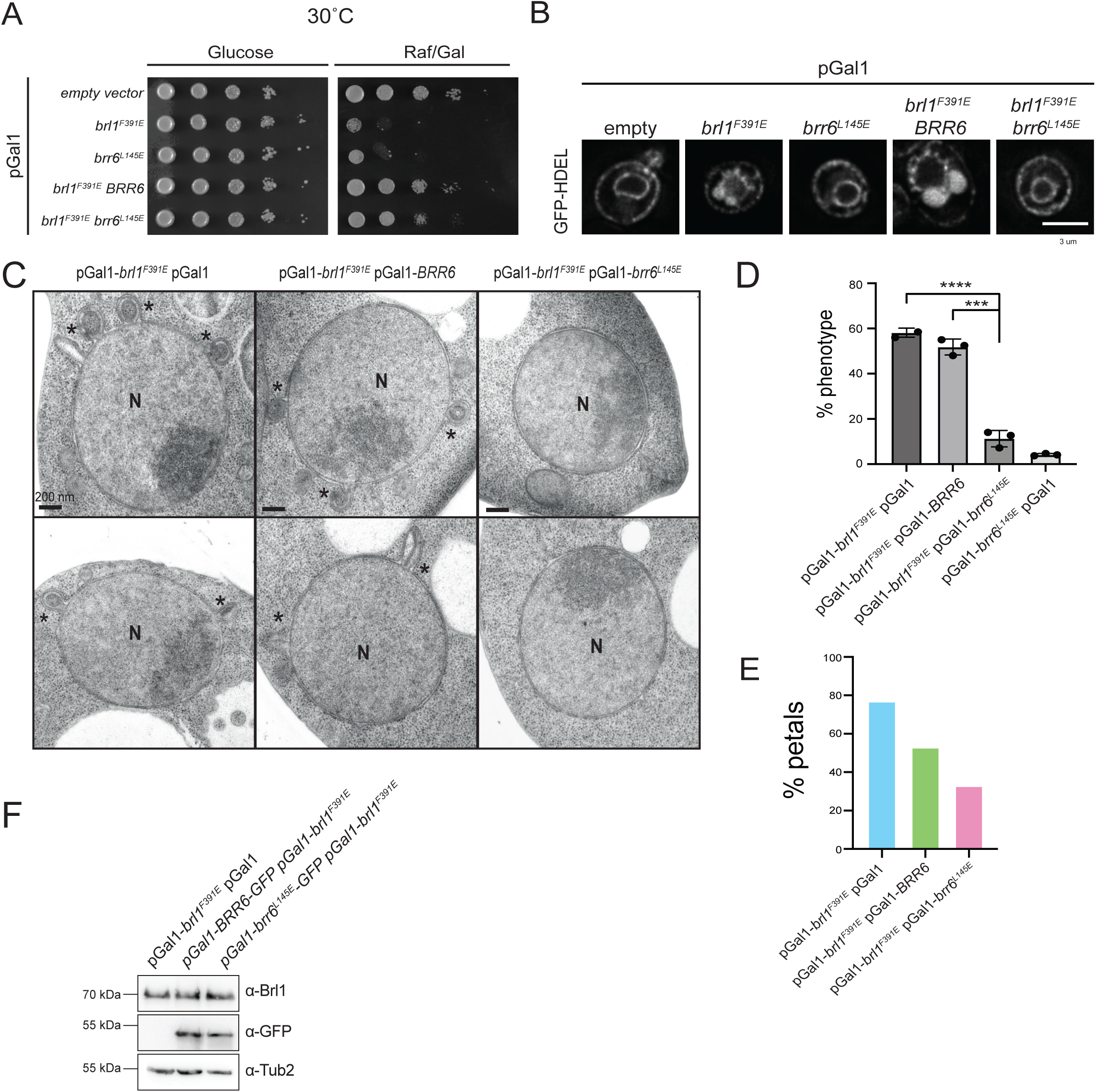
Brl1 and Brr6 AαH domain mutants show cross complementation. **(A)** Overexpression of the *brr6^L145E^*AαH mutant reduces the toxicity of the *brl1^F391E^* AαH mutant. Wild-type yeast cells carrying the indicated plasmids, along with the empty pGal1 plasmid (not shown in the figure), were grown on glucose or galactose/raffinose plates at 30°C. **(B)** Cells expressing the ER marker dsRED-HDEL and the indicated plasmids were imaged by fluorescence microscopy after a 3 hours induction with galactose. Scale bar: 3 µm**. (C)** Electron micrographs of cells overexpressing the indicated plasmids for 3 hours at 30°C. Scale bar: 200 nm. Asterisks indicate petal-like structures at the NE. Abbreviation: N, nucleus. **(D)** Quantification of cells with NE petal-like structures upon overexpression, based on data from (B). Data represent the mean ± SD from three independent experiments (n = 3). Statistical analysis: ANOVA. **(E)** Quantification of the petal-like phenotype observed in (C). Data from a single experiment; the number of cell sections analyzed per strain is indicated. **(F)** Immunoblot showing expression levels of *brl1^F391E^-GFP*, *BRR6-GFP*, and *brr6^L145E^-yeGFP* using the indicated antibodies. Tub2 was used as a loading control.

Consistent with this rescue, analysis of the pGal1-*brl1^F391E^*and pGal1-*brr6^L145E^* phenotype in cells with the NE/ER marker HDEL-eGFP indicated that petal-like structures were only detected in pGal1-*brl1^F391E^* pGal1 and pGal1-*brl1^F391E^* pGal1-*BRR6* cells (Figure 5b, d) while co-expression of pGal1-*brl1^F391E^* and pGal1-*brr6^L145E^* prevented formation of NE petals (Figure 5b, d). This observation was confirmed by EM analysis of pGal1-*brl1^F391E^* pGal1-*brr6^L145E^* cells (Figure 5c, e). We established that expression of Brl1 was similar in all cell types (Figure 5f).

The cross-complementation between Brl1 and Brr6 AαH domain mutants indicates cooperativity between both proteins during early and later steps in NPC assembly.

### Brr6 also functions in INM/ONM fusion during NPC biogenesis

The INM and ONM localization of Brr6 suggests multiple roles of Brr6 in NPC biogenesis (Hodge *et al*., 2010; Zhang *et al*., 2018). To further analyze these functions, we examined six randomly generated conditional-lethal *brr6(ts)* mutants, which carried mutations in the perinuclear space region, including changes to the conserved cysteine residues C96 (*brr6-733*) and C124 (*brr6-19*) (Figure 6a).

**Figure 6.**
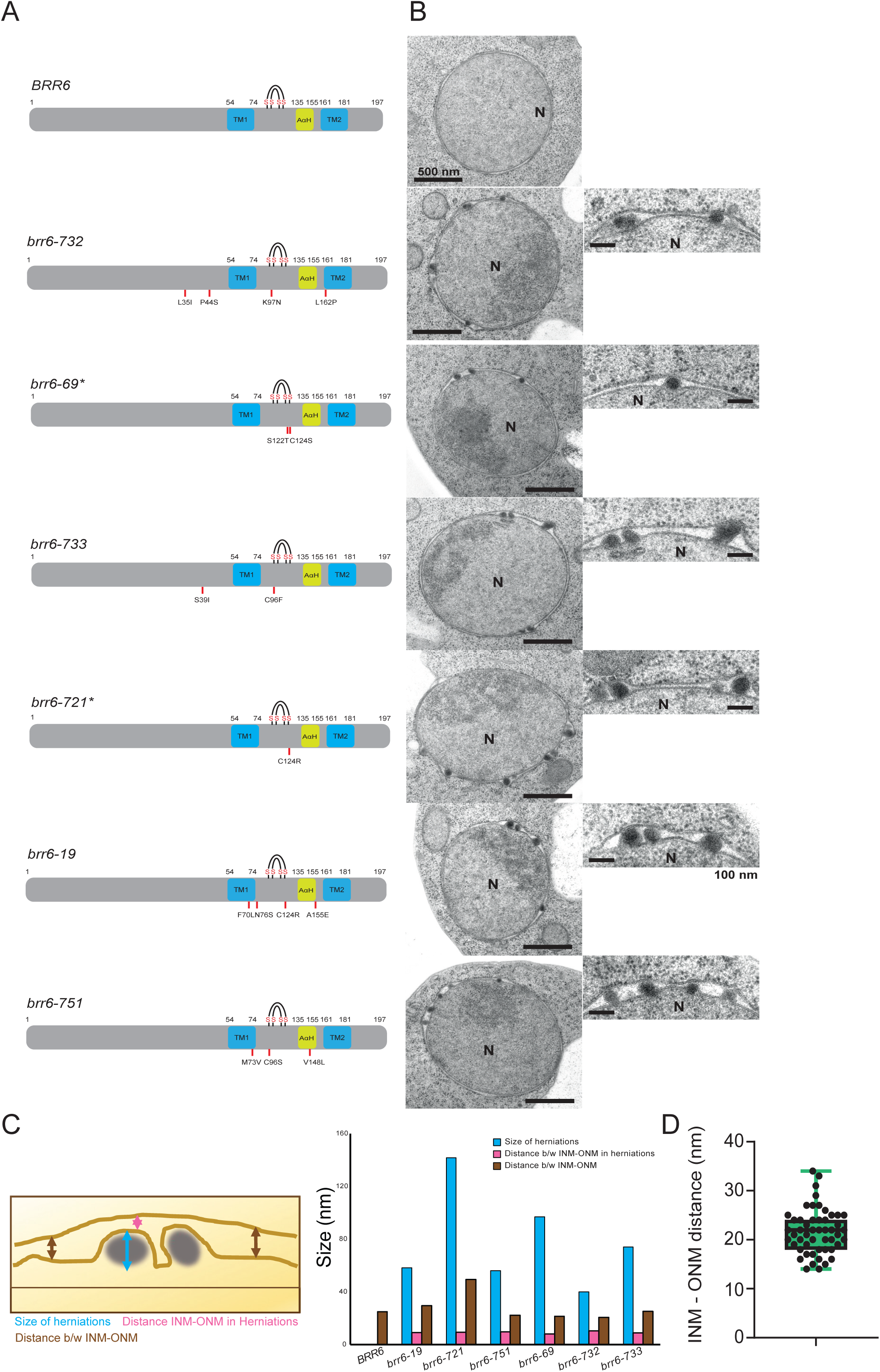
Brr6 also functions in INM/ONM fusion during NPC biogenesis. **(A)** Overview of conditional lethal *brr6(ts)* alleles generated by mutagenic PCR. The resulting *brr6(ts)* alleles were verified by sequencing and integrated into the endogenous *BRR6* locus using a pop-in pop-out strategy. The amino acid substitutions in the various *brr6(ts)* alleles are indicated. TM, AαH, and the four cysteines involved in disulfide bridge formation are also marked. **(B)** The *brr6(ts)* mutants were shifted to 37°C for 3 hours, and their phenotype was analyzed by thin-section EM. All *brr6(ts)* mutants accumulated herniations at the NE. Scale bar: 500 nm; magnified inset: 100 nm. Abbreviation: N, nucleus. **(C)** The schematic on the left illustrates the measured parameters. Quantification of herniation size, the distance between the herniation tip and the ONM, and the distance between the INM/ONM. The indicated parameters were measured from EM sections in (B). 7-15 cell section were analyzed per per strain. **(D)** The distance between the INM and ONM in wild-type cells grown at 30 °C was measured by electron microscopy. Ultrathin sections from 51 cells were analyzed, revealing an average INM–ONM spacing of 22 nm.

For phenotypic analysis, *BRR6* and *brr6(ts)* cells were incubated for 3 hours at the restrictive temperature of 37 °C and examined by EM. All lethal *brr6(ts)* mutants accumulated small to intermediate-sized herniations that distorted the INM (Figure 6b). These herniations, measuring approximately 40-140 nm in size, contacted the ONM without any evidence of membrane fusion (Figure 6c). In addition, the distance between the INM and ONM in wild-type cells incubated at 30 °C was measured to be approximately 22 nm (Figure 6d). In summary, mutations in Brr6 that affect the DHA domain impair INM–ONM fusion during NPC assembly.

### The N-terminus of Brl1 fulfills an essential function and interacts with the Nic96 complex

Brl1 possesses a relatively long N-terminal extension of 299 amino acids, which— based on its topology—is exposed to the cytoplasm or nucleoplasm (Zhang *et al*., 2018). In contrast, its C-terminal extension comprises only 42 amino acids. Interestingly, the N-terminus of *S. cerevisiae* Brl1 shares sequence homology with the N-terminus of *S. pombe* Brr6 (Supplementary Figure 2).

To assess the functional relevance of these regions, we replaced them with the corresponding segments from Brr6 (53 aa N-terminal / 15 aa C-terminal). While the *N-BRR6–BRL1* fusion was non-functional (Figure 7b, row 4), the *BRL1–C-BRR6* chimera retained the essential function of *BRL1* (Figure 7b, row 5), indicating that the N-terminus of Brl1 carries a critical function. Similarly, analysis of *BRR6* function revealed that the N-terminal Brl1–Brr6 fusion was non-functional (Figure 7C, row 6), whereas the C-terminal Brr6–Brl1 fusion retained functionality (Figure 7C, row 7).

**Figure 7.**
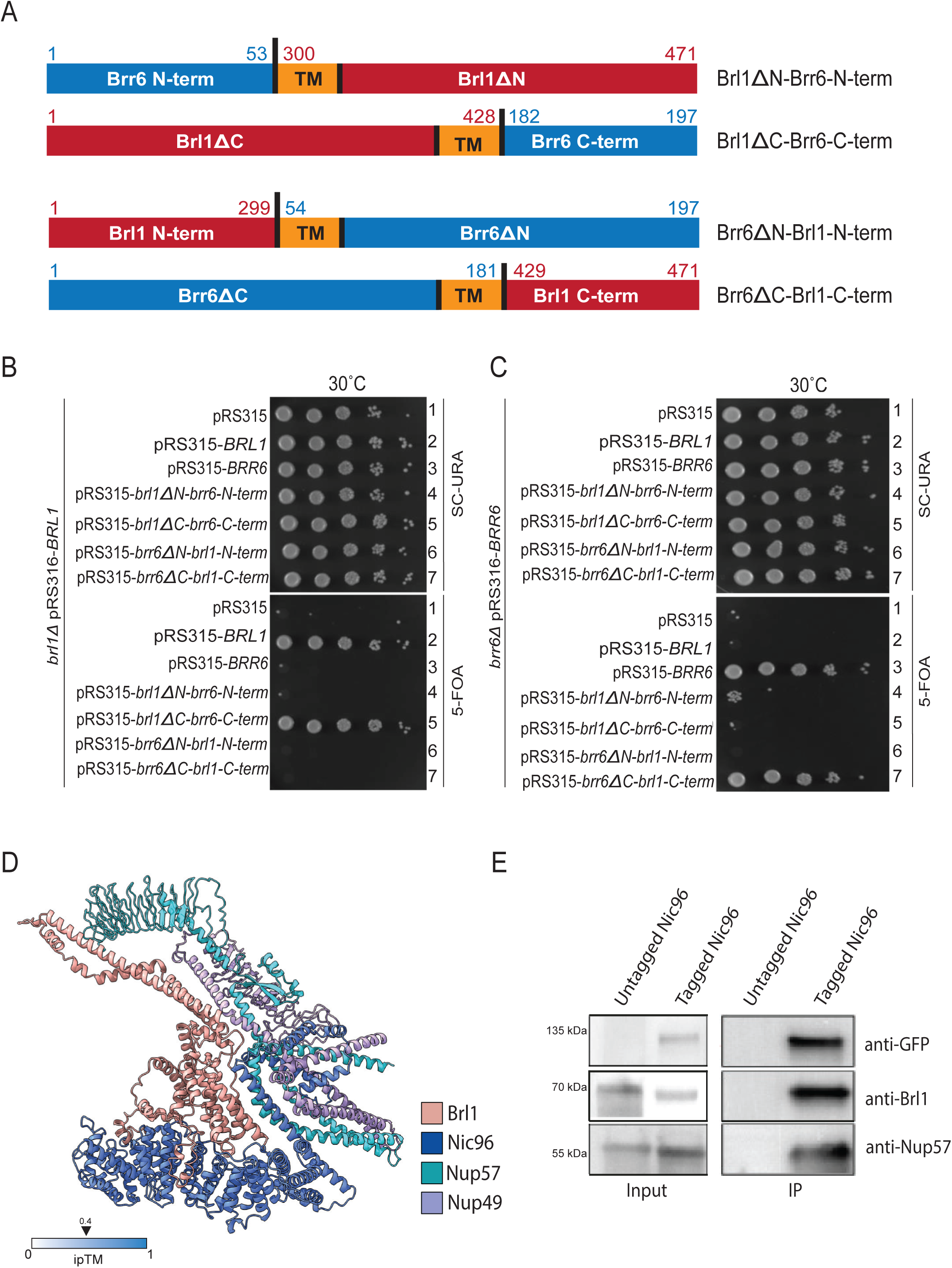
The N-terminus of Brl1 fulfills an essential function and interacts with the Nic96 complex. **(A)** Schematic representation of Brl1-Brr6 fusion proteins and their nomenclature. The amino acid numbers at the fusion junctions are indicated. The cartoon also depicts the TM domains of Brl1 and Brr6. **(B, C)** *BRL1*, *BRR6*, and *BRL1-BRR6* fusion constructs were expressed from their respective endogenous promoters using the *LEU2*-based pRS315 plasmid and transformed into *BRL1* **(B)** or *BRR6* **(C)** shuffle strains. Transformants were grown at 30 °C on SC–URA and 5-FOA plates. Growth on 5-FOA indicates functional complementation. The *brl1ΔC-brr6-C-term* construct was functional for Brl1, while the fusion containing Brl1 with the N-terminus of Brr6 (*brl1ΔN-brr6-N-term*) was non-functional (B, rows 5 and 4, respectively) indicating an essential role of the N-terminus of Brl1. Similarly, *brr6ΔN-brl1-N-term* failed to complement *BRR6* function, while *brr6ΔC-brl1-C-term* was functional (C, rows 6 and 7). **(D)** AlphaFold prediction showing a potential interaction between the N-terminus of Brl1 and nucleoporins Nic96, Nup57, and Nup49. **(E)** Co-immunoprecipitation of Brl1 with Nic96 and Nup57. Cells expressing either *NIC96* or *NIC96-GFP*, both with *BRL1-HA*, were lysed, and Nic96-GFP was immunoprecipitated using GFP nanobody beads. Eluted proteins were analyzed by immunoblotting with the indicated antibodies. Brl1 and Nup57 were specifically enriched in the Nic96-GFP pulldown.

AlphaFold predictions revealed a potential interaction between the N-terminal region of Brl1 and Nic96, which forms a complex with Nup57 (Figure 7d). We validated this prediction by demonstrating co-immunoprecipitation between Brl1, Nic96, and Nup57 (Figure 7e). These findings suggest that the interaction between Nic96 and the N-terminus of Brl1 is likely critical for Brl1’s role in NPC assembly.

### Brl1 and Brr6 promote NE fusion

It has been proposed that Brl1 and Brr6 promote fusion of the INM and ONM during NPC assembly by interacting head-to-head via the tips of their respective DAH domains (Figure 1d). However, AlphaFold analysis of Brl1 and Brr6 suggests that the lengths of their DAH domains, even in head-to-head configurations, are with 16 nm insufficient to span the ∼22 nm width of the perinuclear space in yeast (Figure 6d). Notably, as the INM becomes deformed by the deposition of Nups from the nuclear side, the local narrowing of the perinuclear space may bring the DAH domains of Brl1 at the INM and Brr6 at the ONM into close proximity, enabling their interaction. Based on this model, we hypothesized that artificially extending the DAH domains beyond a critical distance could allow Brl1 and Brr6 to interact and promote membrane fusion independently of INM deformation.

To test this, we extended the DAH domains of Brl1 and Brr6 by inserting 21 amino acids into Brl1 (Brl1^ePNS^) and 15 into Brr6 (Brr6^ePNS^), increasing their predicted lengths from 9 nm to 16 nm and from 7 nm to 12 nm, respectively (Figure 8a). Both, the *brl1^ePNS^* and *brr6^ePNS^*constructs were non-functional in yeast (Figure 8b, c). Strikingly, expression of these extended variants *brl1^ePNS^* and *brr6^ePNS^* resulted in a pronounced toxic phenotype, which was not observed upon overexpression of the wild-type *BRL1* or *BRR6* (Figure 8d, e).

**Figure 8.**
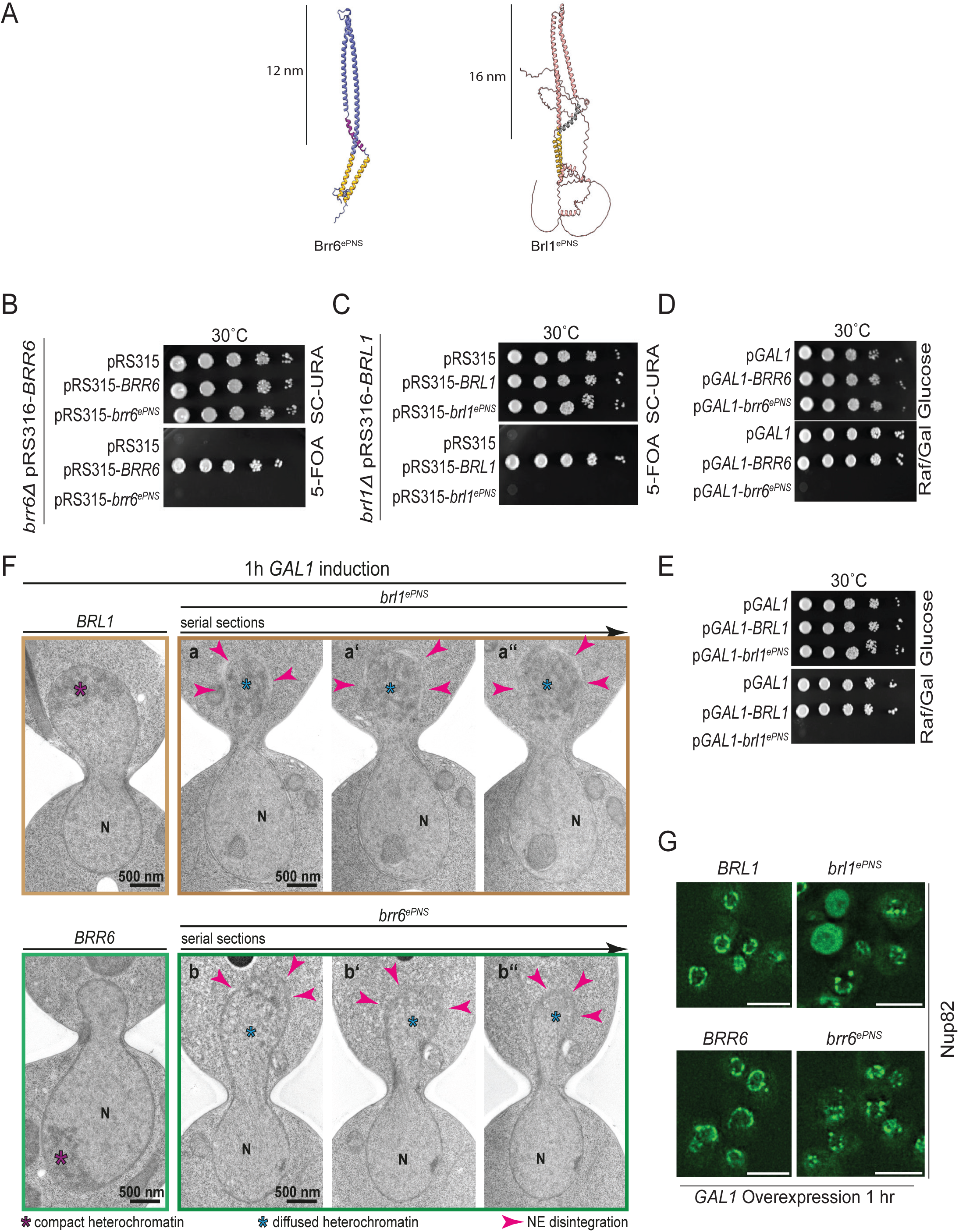
Brl1 and Brr6 promote NE fusion. **(A)** AlphaFold structural predictions of the extended DHA regions of Brl1 (Brl1^ePNS^) and Brr6 (Brr6^ePNS^). **(B, C)** *brl1^ePNS^* and *brr6^ePNS^* constructs are non-functional in a plasmid shuffle assay at 30°C, whereas wild-type *BRL1* and *BRR6* are functional. **(D, E)** Overexpression of pGal1-*Brl1^ePNS^* and pGal1-*Brr6^ePNS^* is toxic to cells, while overexpression of wild-type *BRL1* and *BRR6* does not affect cell growth. **(F)** EM analysis of cells expressing pGal1-*BRL1^ePNS^* and pGal1-*BRR6^ePNS^* after 60 minutes of galactose induction. Panels a–a** and b–b** show 80 nm serial sections of the same cell. Arrowheads indicate regions of the NE undergoing disintegration. Purple asterisks mark compact chromatin; blue asterisks denote decondensed chromatin adjacent to disintegrating NE regions. Abbreviation: N, nucleus. Scale bar: 500 nm. **(G)** pGal1-*BRL1^ePNS^* and pGal1-*BRR6^ePNS^* were expressed in *NUP82-GFP* cells following 1 hour galactose induction. Scale bars: 5 µm.

To understand the cause of the lethality associated with *brl1^ePNS^*and *brr6^ePNS^*, we analyzed *NUP82*-GFP–tagged cells expressing either the wild-type or extended versions of *BRL1* and *BRR6* under the control of the pGAL1 promoter. Fluorescence microscopy of cells 60 minutes after pGal1 induction revealed a pronounced defect in NPC organization (Figure 8g). Some cells exhibited a uniform Nup82-GFP signal distributed throughout the cytoplasm, indicating severe disruption of the NE and NPC structures (Figure 8g). These findings indicate that expression of the extended versions of *BRL1* and *BRR6* compromises nuclear NE NPC integrity.

To gain deeper insight into the defects observed by fluorescence microscopy data, we performed EM analysis. Thin-serial section EM of *brl1^ePNS^*- and *brr6^ePNS^*-expressing cells revealed NE abnormalities, including cells with partially disintegrated NEs visible across multiple serial sections (Figure 8f). Notably, such defects were absent in pGal1-*BRL1* and pGal1-*BRR6* control cells.

Taken together, the toxic phenotype of *brl1^ePNS^* and *brr6^ePNS^* mutant cells, the disrupted NE/NPC organization observed by fluorescence microscopy, and the NE disintegration detected by EM strongly support a model in which spatial regulation of DAH domain interactions in Brl1 and Brr6 is essential for controlled INM–ONM fusion during NPC assembly.

## Discussion

The inside-out NPC assembly pathway involves the fusion of the INM and ONM (Otsuka & Ellenberg, 2018), allowing the NPCs to become embedded in the nascent openings of the NE. Despite its importance, the mechanism underlying this membrane fusion has remained elusive. However, the discovery of Brl1 and Brr6 in yeast, two paralogues interacting, integral membrane proteins of the NE, has shed light on this process (de Bruyn Kops & Guthrie, 2001; Hodge *et al*., 2010; Kralt *et al*., 2022; Lo Presti *et al*, 2007; Lone *et al*., 2015; Saitoh *et al*, 2005; Vitale *et al*., 2022; Zhang *et al*., 2018). These proteins are essential for nuclear transport, specifically localize to NPC assembly sites and not to fully assembled NPCs, and facilitate the fusion of the INM and ONM during NPC assembly, suggesting they are components of the membrane fusion machinery required for NPC integration into the NE. Structural predictions of Brl1 and Brr6 revealed a DHA stabilized by two disulfide bonds near its tip, along with an AαH at the base of the DHA (Kralt *et al*., 2022; Vitale *et al*., 2022). These features suggest the presence of two structural elements that may contribute to the INM/ONM fusion process during NPC assembly. In addition, Brl1 and Brr6 contain N-terminal domains, the functional relevance of which has not yet been analyzed.

Previous studies have shown that overexpression of *brl1* mutants with a defective AαH results in excessive proliferation of the INM, forming large petal-like structures that deform the ONM but do not lead to membrane fusion (Kralt *et al*., 2022; Vitale *et al*., 2022). Mutations that disrupt the integrity of the DHA cause a temperature-sensitive phenotype, characterized by the accumulation of herniations, which is an indication of failed INM/ONM fusion (Figure 6) (de Bruyn Kops & Guthrie, 2001; Saitoh *et al*., 2005; Zhang *et al*., 2018). Together, these findings suggest that DHA and AαH both have functions late in NPC assembly, in the fusion of the INM/ONM.

We observed distinct phenotypes in *brr6* mutants affecting the AαH and the DHA, consistent with roles in both early and late stages of NPC assembly. Overexpression of *BRR6* AαH mutants disrupts the NE localization of most Nups (Figures 2). A similar phenotype on Nup82 was observed in the cold-sensitive *brr6^L145E^* AαH mutant. Additionally, the number of NPCs drastically decreased during incubation of *brr6^L145E^* cells at 16°C, indicating an NPC assembly defect without the formation of herniations (Figure 4e). This phenotype raises the question of why it was not observed in earlier studies, such as those employing Brr6 degron or Brl1/Brr6 double degron systems, which reported the accumulation of NE herniations upon protein degradation (Zhang *et al*., 2018). One possibility is that the early function of Brr6 in NPC assembly requires only low protein levels, and thus is less sensitive to depletion than its later roles. Additionally, mutations in the DHA region of *BRR6* led to the formation of herniations consistent with previous observations (de Bruyn Kops & Guthrie, 2001) (Figure 6), indicating that the primary function of the DHA in Brr6 is facilitating INM/ONM fusion during NPC assembly.

The cross-complementation observed between the *brl1* and *brr6* AαH mutants (Figure 5) suggests that this early function of Brr6 is carried out in coordination with its interacting partner Brl1, likely at the INM, where both proteins probably engage via their AαH domains (Figure 1) (Zhang *et al*., 2018). Screening *BRL1* for cold-sensitive alleles, particularly within the region encoding the N-terminus (see below), may identify mutants that disrupt its early role in NPC assembly.

The ability of the Brl1 N-terminus to bind the Nic96 complex (Figure 7), an essential component in the early stages of NPC assembly (Onischenko *et al*., 2020), along with the similar NPC assembly defects observed in *brr6^L145E^* and *nic96(ts)* mutant cells and the genetic interaction between *brr6-1* (Arg110Lys) with the Δ*nic96(532-839)* allele (de Bruyn Kops & Guthrie, 2001; Grandi *et al*, 1995; Zabel *et al*, 1996), indicates that the early function of the Brl1–Brr6 complex involves interaction with Nic96. Because Nic96 is mislocalized in cells expressing pGal1*-brr6^L145E^* and pGal1*-brr6^F152E^* (Figure 3b), we propose that the Brl1–Brr6–Nic96 interaction is essential for the proper recruitment of the Nic96 complex to early NPC assembly sites. Brl1–Brr6 may function as a scaffold that facilitates the incorporation of the Nic96 complex into NPC assembly intermediates.

As more Nups are deposited at these early NPC assembly sites, the INM begins to deform inward into the perinuclear space region. This membrane deformation reduces the local distance between the INM and ONM from approximately 22 nm to around 15 nm (Figure 6). We propose that this decrease in distance is sufficient to allow a head-to-head interaction between the DHA domains of Brl1 on the INM and Brr6 on the ONM, which is based on AlphaFold predications close to 16 nm (Figure 1). Support for this spatially regulated interaction model comes from the phenotype of *brl1* and *brr6* mutants in which the DHA was artificially extended, from 9 nm to 16 nm for Brl1 and from 7 nm to 12 nm for Brr6. Thus, head-to-head DAH interactions between Brl1^ePNS^–Brr6 and Brl1–Brr6^ePNS^ likely exceed the critical ∼22 nm distance between the INM and ONM.

These extended DHA mutants were non-functional in a shuffle strain, but their overexpression caused a severe toxic growth defect (Figure 8). Further analysis revealed defective NPC distribution along the NE and, by EM, evidence of NE disintegration. This phenotype is likely caused by uncontrolled fusion between the INM and ONM, possibly driven by unregulated interactions between the extended DHA head domains of Brl1 and Brr6 on opposite sides of the NE. While the DHA domain likely mediates this bridging function between Brl1 and Brr6 as indicated by the phenotype of conditional lethal DHA mutants and the spatial distribution of both proteins on the INM and ONM (de Bruyn Kops & Guthrie, 2001; Zhang *et al*., 2018), the AαH of Brl1 may promote INM–ONM fusion once the two membranes are in close proximity. How Apq12 cooperates with Brl1 and Brr6 during INM/ONM fusion in NPC assembly remains to be elucidated in future studies (Hodge *et al*., 2010; Lone *et al*., 2015; Scarcelli *et al*., 2007; Zhang *et al*., 2021).

Brl1 and Brr6, which are stabilized by disulfide bonds in the DHA domain, are found only in organisms that retain the NE during mitosis (Tamm *et al*, 2011). In many organisms, such as *Schizosaccharomyces pombe*, Brl1/Brr6 functions are carried out by a single homologue, Brr6 (Tamm *et al*., 2011). In these cases, the solitary Brr6 protein likely needs to engage in both lateral and head-to-head interactions, as observed for Brl1 and Brr6 in budding yeast. Interestingly, a remote homology search identified CLCC as a putative Brl1/Brr6 homologue with potential roles in NPC assembly (Alyssa J. Mathiowetz *et al*, 2024). It will be intriguing to determine which functional domains within CLCC mediate Nup recruitment and INM/ONM fusion.

## Acknowledgment

This work was funded by the Deutsche Forschungsgemeinschaft (SFB1638). We thank Dr. Iain Hagan, University of Manchester, for kindly providing the conditional lethal *brr6(ts)* cells and the Brl1 antibodies. We thank Ed Hurt and Martin Beck for the anti-Nup57 antibody. We thank the EM core facility of Heidelberg University (EMCF) for their technical support and Uta Hasselmann for assistance.

## Autor contributions

ES and SM have written the manuscript with the help from AN. All EM analysis and the quantifications associated with it was done by AN including Figure 2, 3, 5, 6 and 8. SM has performed experiments described in Figures 1, 4, 7 and 8. AK has performed experiments described in Figures 2, 3, 5.

## Supplementary Figures

**Figure S1. Analysis of *brr6^L145E^* and *brr6^F152E^* overexpression phenotypes. Extension of Figure 3**.

**(A)** Schematic representation of the NPC with color-coded components: transmembrane ring (green), central FG-Nups (purple), nuclear basket Nups (red), linker Nups (blue), inner ring Nups (ochre), and outer ring Nups (light blue).

**(B)** Overexpression of *brr6^F152E^* alters NE localization of Nsp1 and Nup159, whereas Nup2 and Pom152 are only minimally affected. Line scans on the right show NE distribution of the indicated Nups in cells carrying either empty pGal1 or pGal1-*brr6^F152E^* plasmids. Cells were incubated for 3 hours in raffinose/galactose medium to induce expression from the pGal1 promoter. Scale bar: 5 µm. **(C)** As in (B), using the indicated yeast strains expressing tdTomato-tagged Nup constructs. Cells additionally carried pGal1, pGal1-*BRR6*, pGal1-*brr6^L145E^*, or pGal1-*brr6^F152E^*, which were overexpressed for 3 hours. Scale bar: 5 µm.

**Figure S2.** Comparison of *S. cerevisiae* Brl1, Brr6 and *S. pombe* Brr6.

Amino acid sequence alignment of Brl1 and Brr6 from *Saccharomyces cerevisiae* and Brr6 from *Schizosaccharomyces pombe*. The alignment highlights conservation not only within the DHA domain containing four cysteine residues, but also between the N-terminal regions of *S. cerevisiae* Brl1 *and S. pombe* Brr6.

## MATERIALS AND METHODS

### Yeast strains and plasmids

Yeast strains used in this study are derivatives of ESM356-1 (MATa *ura3-52 trp1Δ63 his3Δ200 leu2Δ1*). Endogenous gene tagging and gene deletions were performed using PCR-based integration methods as described by (Janke *et al*., 2004). Strains were cultured in YPD (yeast extract, peptone, and glucose), SC (synthetic complete) medium, or SC selective medium lacking specific amino acids or bases, as indicated. For induction of proteins under control of the pGal1 promoter, cells were grown in SC medium containing raffinose and supplemented with 2% galactose. Growth assays were performed by growing cells overnight in selective medium, adjusting cultures to an OD_600_ of 1.0, and spotting 10-fold serial dilutions onto selective agar plates. Plates were incubated at the specified temperatures. For protein analysis by immunoblotting, yeast extracts were prepared using alkaline lysis followed by TCA precipitation (Knop *et al*, 1999).

### Electron microscopy

High-pressure frozen yeast samples were processed for EM analysis as described in the following: cells were collected onto a 0.45 μm polycarbonate filter (Millipore) using vacuum filtration and subsequently frozen using a high-pressure freezing device (HPM010, Abra-Fluid, Switzerland). Freeze substitution was performed in an EM-AFS2 system (Leica Microsystems, Vienna, Austria) using a solution containing 0.2% uranyl acetate and 1% water in anhydrous acetone. Samples were stepwise infiltrated with Lowicryl HM20 resin (Polysciences, Inc., Warrington, PA) starting at −90 °C. Polymerization was carried out upon UV light for 48 hours at −45 °C, followed by gradual warming to 20 °C.

In resin embedded yeast cells were serial sectioned at a thickness of 80 nm using a Reichert Ultracut S microtome (Leica Instruments, Vienna, Austria) and fished on a slot grids, covered with a thin plastic foil. Sections were post-stained with 3% uranyl acetate and lead citrate and imaged using a JEOL JEM-1400 transmission electron microscope (JEOL Ltd., Tokyo, Japan), operated at 80 kV and equipped with a 4k × 4k digital camera (F416, TVIPS, Gauting, Germany). Image brightness and contrast were adjusted using Fiji (NIH, Bethesda, MD).

For immunogold labeling, primary antibody against Nsp1 was used. Serial sections were mounted on slot grids, incubated with blocking buffer (1.5% BSA, 0.1% fish skin gelatin in phosphate-buffered saline [PBS]), followed by incubation with the primary antibody (rabbit anti-Nsp1). After washing in PBS, sections were incubated with a secondary linker antibody (rabbit anti-mouse), and labeled using protein A–gold conjugates (15 nm; Utrecht University, Utrecht, Netherlands). Post-staining was performed as above using 3% uranyl acetate and lead citrate.

### Immunoprecipitation

Cells equivalent to 25 OD_600_ units were harvested and resuspended in lysis buffer (20 mM Tris–Cl, pH 8.0; 150 mM NaCl; 5 mM MgCl_2_; 10% glycerol) supplemented with 10 mM NaF, 60 mM β-glycerophosphate, 1 mM PMSF, and 1 tablet of EDTA-free protease inhibitor cocktail (Roche) per 50 ml. Cell lysis was performed using a FastPrep homogenizer (MP Biomedicals) with the addition of glass beads (BioSpec Products). After lysis, Triton X-100 was added to a final concentration of 0.5%, and samples were incubated on ice for 10 minutes.

The lysates were clarified by centrifugation to separate soluble proteins from debris. The supernatant was incubated with GFP-Trap agarose beads (Chromotek) for 2 hours at 4 °C with gentle rotation. Beads were then washed three times with lysis buffer containing 0.1% Triton X-100 and twice with wash buffer (20 mM Tris–Cl, pH 8.0; 150 mM NaCl; 5 mM MgCl_2_). Bound proteins were eluted by boiling the beads in 50 μl of 2× Laemmli sample buffer for 5 minutes at 95 °C and analyzed by SDS-PAGE and immunoblotting.

### Fluorescence microscopy

Cell imaging was performed using a DeltaVision RT system (Applied Precision Ltd.) based on an Olympus IX71 microscope and equipped with a Photometrics CoolSnap HQ camera (Roper Scientific), a 100×/1.4 NA Super-Plan Apochromat oil immersion objective (Olympus), and a four-color Standard Insight SSI module light source. The system was controlled via a workstation running the CentOS operating system and softWoRx software (Applied Precision Ltd.).

Images were acquired using either the GFP or mCherry channels under identical exposure times and illumination settings to enable direct comparison across samples. A single optical z-stack was collected per field of view and used for analysis. Image deconvolution was performed using softWoRx, and further image processing was conducted with ImageJ (National Institutes of Health, Bethesda, MD).

Most imaging experiments and quantitative analyses were independently repeated three times. Data analysis and statistical evaluation were performed using GraphPad Prism software.

### Antibodies

The antibodies of this study were used as follow: mouse anti-Nsp1 (immuno-EM, 1:100; ab4641; Abcam), rabbit anti-Tub2 (immunoblot, 1:1000; made in-house), rabbit anti-Brl1 (gift from Dr. I. Hagan, Manchester; 1:500) and rabbit anti-Nup57 antibodies (gift from Ed Hurt, Heidelberg; 1:100).

## References

Alyssa J. Mathiowetz, Emily S. Meymand, Kirandeep K. Deol, Güneş Parlakgül, Mike Lange, Stephany P. Pang, Melissa A. Roberts, Emily F. Torres, anielle M. Jorgens, Reena Zalpuri et al (2024) CLCC1 promotes hepatic neutral lipid flux and nuclear pore complex assembly. bioRxiv

Beck M, Hurt E (2017) The nuclear pore complex: understanding its function through structural insight. Nat Rev Mol Cell Biol 18: 73–89

de Bruyn Kops A, Guthrie C (2001) An essential nuclear envelope integral membrane protein, Brr6p, required for nuclear transport. EMBO J 20: 4183–4193

Frey S, Gorlich D (2007) A saturated FG-repeat hydrogel can reproduce the permeability properties of nuclear pore complexes. Cell 130: 512–523

Grandi P, Schlaich N, Tekotte H, Hurt EC (1995) Functional interaction of Nic96p with a core nucleoporin complex consisting of Nsp1p, Nup49p and a novel protein Nup57p. EMBO J 14: 76–87

Hodge CA, Choudhary V, Wolyniak MJ, Scarcelli JJ, Schneiter R, Cole CN (2010) Integral membrane proteins Brr6 and Apq12 link assembly of the nuclear pore complex to lipid homeostasis in the endoplasmic reticulum. J Cell Sci 123: 141–151

Janke C, Magiera MM, Rathfelder N, Taxis C, Reber S, Maekawa H, Moreno-Borchart A, Doenges G, Schwob E, Schiebel E et al (2004) A versatile toolbox for PCR-based tagging of yeast genes: new fluorescent proteins, more markers and promoter substitution cassettes. Yeast 21: 947–962

Jumper J, Evans R, Pritzel A, Green T, Figurnov M, Ronneberger O, Tunyasuvunakool K, Bates R, Zidek A, Potapenko A et al (2021) Highly accurate protein structure prediction with AlphaFold. Nature 596: 583-+

Knop M, Siegers K, Pereira G, Zachariae W, Winsor B, Nasmyth K, Schiebel E (1999) Epitope tagging of yeast genes using a PCR-based strategy: More tags and improved practical routines. Yeast 15: 963–972

Kralt A, Wojtynek M, Fischer JS, Agote-Aran A, Mancini R, Dultz E, Noor E, Uliana F, Tatarek-Nossol M, Antonin W et al (2022) An amphipathic helix in Brl1 is required for nuclear pore complex biogenesis in S. cerevisiae. Elife 11

Lin DH, Hoelz A (2019) The Structure of the Nuclear Pore Complex (An Update). Annu Rev Biochem 88: 725–783

Lo Presti L, Cockell M, Cerutti L, Simanis V, Hauser PM (2007) Functional characterization of Pneumocystis carinii brl1 by transspecies complementation analysis. Eukaryotic Cell 6: 2448–2452

Lone MA, Atkinson AE, Hodge CA, Cottier S, Martinez-Montanes F, Maithel S, Mene-Saffrane L, Cole CN, Schneiter R (2015) Yeast Integral Membrane Proteins Apq12, Brl1, and Brr6 Form a Complex Important for Regulation of Membrane Homeostasis and Nuclear Pore Complex Biogenesis. Eukaryot Cell 14: 1217–1227

Munro S, Pelham HR (1987) A C-terminal signal prevents secretion of luminal ER proteins. Cell 48: 899–907

Onischenko E, Noor E, Fischer JS, Gillet L, Wojtynek M, Vallotton P, Weis K (2020) Maturation Kinetics of a Multiprotein Complex Revealed by Metabolic Labeling. Cell 183: 1785-+

Onischenko E, Tang JH, Andersen KR, Knockenhauer KE, Vallotton P, Derrer CP, Kralt A, Mugler CF, Chan LY, Schwartz TU et al (2017) Natively Unfolded FG Repeats Stabilize the Structure of the Nuclear Pore Complex. Cell 171: 904–917 e919

Otsuka S, Bui KH, Schorb M, Hossain MJ, Politi AZ, Koch B, Eltsov M, Beck M, Ellenberg J (2016) Nuclear pore assembly proceeds by an inside-out extrusion of the nuclear envelope. Elife 5

Otsuka S, Ellenberg J (2018) Mechanisms of nuclear pore complex assembly - two different ways of building one molecular machine. FEBS Lett 592: 475–488

Otsuka S, Steyer AM, Schorb M, Heriche JK, Hossain MJ, Sethi S, Kueblbeck M, Schwab Y, Beck M, Ellenberg J (2018) Postmitotic nuclear pore assembly proceeds by radial dilation of small membrane openings. Nat Struct Mol Biol 25: 21–28

Saitoh Y, Ogawa K, Nishimoto T (2005) Brl1p - A novel nuclear envelope protein required for nuclear transport. Traffic 6: 502–517

Scarcelli JJ, Hodge CA, Cole CN (2007) The yeast integral membrane protein Apq12 potentially links membrane dynamics to assembly of nuclear pore complexes. J Cell Biol 178: 799–812

Strambio-De-Castillia C, Niepel M, Rout MP (2010) The nuclear pore complex: bridging nuclear transport and gene regulation. Nat Rev Mol Cell Biol 11: 490–501

Tamm T, Grallert A, Grossman EP, Alvarez-Tabares I, Stevens FE, Hagan IM (2011) Brr6 drives the *Schizosaccharomyces pombe* spindle pole body nuclear envelope insertion/extrusion cycle. J Cell Biol 195: 467–484

Thaller DJ, Lusk CP (2018) Fantastic nuclear envelope herniations and where to find them. Biochemical Society Transactions 46: 877–889

Vitale J, Khan A, Neuner A, Schiebel E (2022) A perinuclear alpha-helix with amphipathic features in Brl1 promotes NPC assembly. Mol Biol Cell 33: ar35

Wente SR, Blobel G (1993) A temperature-sensitive NUP116 null mutant forms a nuclear envelope seal over the yeast nuclear pore complex thereby blocking nucleocytoplasmic traffic. J Cell Biol 123: 275–284

Winey M, Yarar D, Giddings TH, Jr., Mastronarde DN (1997) Nuclear pore complex number and distribution throughout the Saccharomyces cerevisiae cell cycle by three-dimensional reconstruction from electron micrographs of nuclear envelopes. Mol Biol Cell 8: 2119–2132

Zabel U, Doye V, Tekotte H, Wepf R, Grandi P, Hurt EC (1996) Nic96p is required for nuclear pore formation and functionally interacts with a novel nucleoporin, Nup188p. J Cell Biol 133: 1141–1152

Zhang W, Khan A, Neuner A, Vitale J, Rink K, Lüchtenborg C, Brügger B, Söllner TH, Schiebel E (2021) A short perinuclear amphipathic α-helix in Apq12 promotes nuclear pore complex biogenesis. Open biology doi: 10.1098/rsob.210250.

Zhang W, Neuner A, Ruthnick D, Sachsenheimer T, Luchtenborg C, Brugger B, Schiebel E (2018) Brr6 and Brl1 locate to nuclear pore complex assembly sites to promote their biogenesis. J Cell Biol 217: 877–894

