## Supplementary figures and images for "Multifunctional Roles of Brr6 and Brl1 in Nuclear Envelope Fusion During Nuclear Pore Complex Biogenesis"

### Supplemental Figures 1 and 2

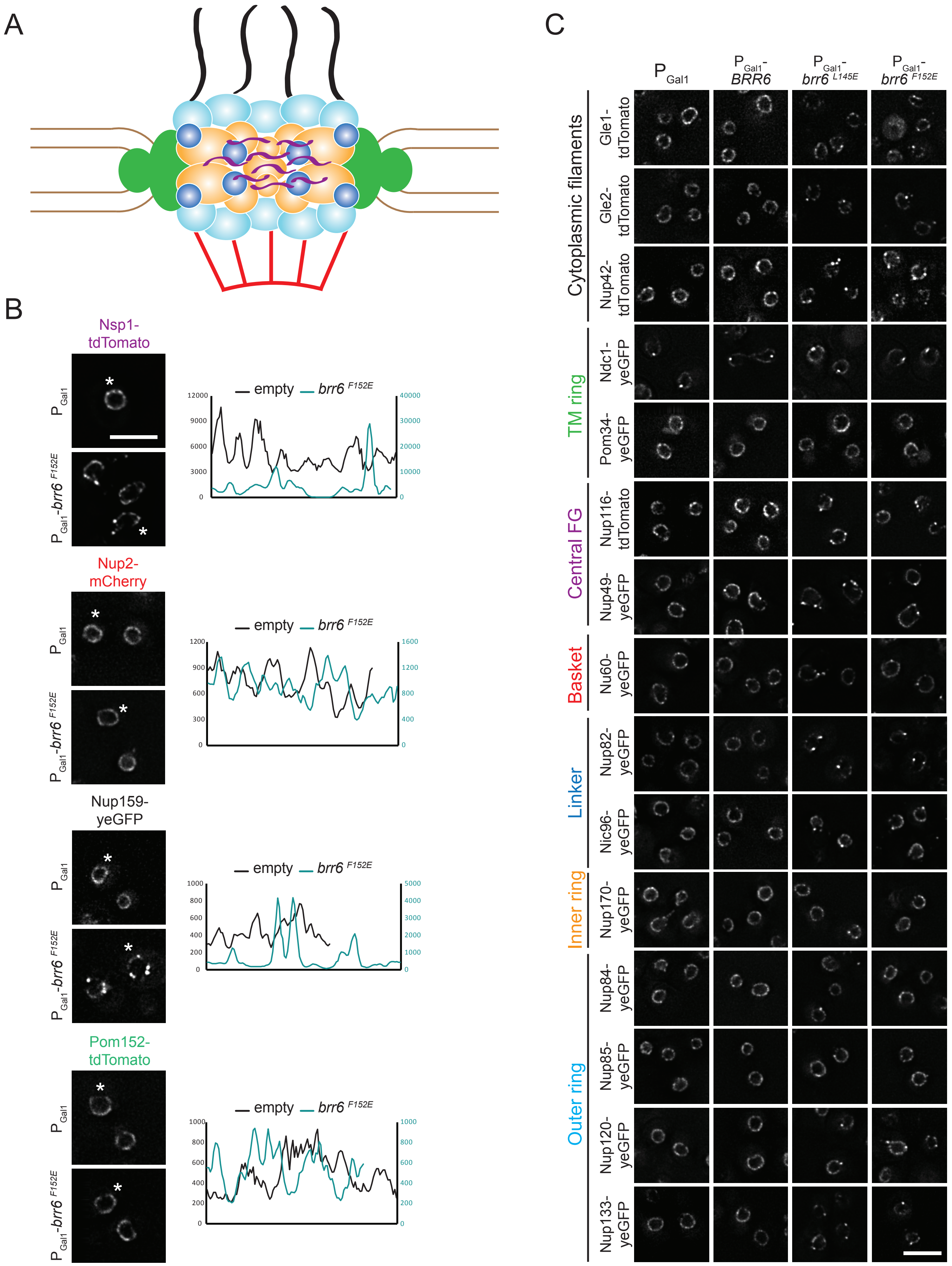

#### Supplementary Figure 2

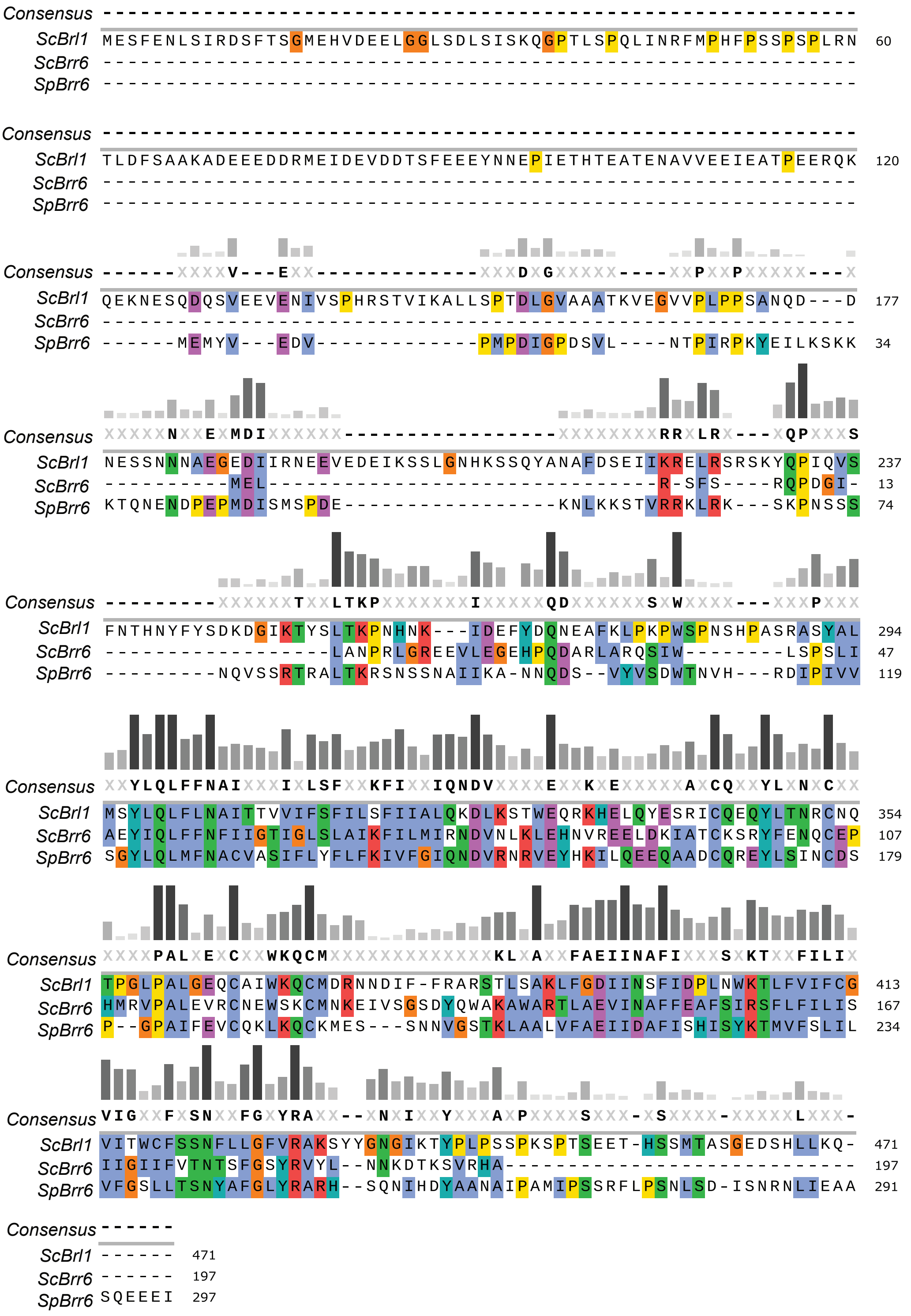
